# Separating sensory from timing processes: a cognitive encoding and neural decoding approach

**DOI:** 10.1101/2024.06.24.600536

**Authors:** Christina Yi Jin, Anna Razafindrahaba, Raphaël Bordas, Virginie van Wassenhove

## Abstract

The internal clock is a psychological model for timing behavior. According to information theory, psychological time might be a manifestation of information flow during sensory processing. Herein, we tested three hypotheses: (1) whether sensory adaptation reduces (or novelty increases) the rate of the internal clock (2) whether the speed of the clock reflects the amount of cortical sensory processing? (3) whether motion tunes clock speed.

The current study used an oddball paradigm in which participants detected duration changes while being recorded with electroencephalography (EEG). For data analysis, we combined cognitive modeling with neural decoding techniques. Specifically, we designed Adaptive-Thought-of-Control (ACT-R) models to explain human data and linked them to the sensory EEG features discovered through machine learning.

Our results indicate that timing performance is influenced by both timing and non-timing factors. The internal clock may reflect the amount of sensory processing, thereby clarifying a long-standing sensory timing mystery.

## Introduction

Time perception is fundamental to human cognition, affecting various cognitive functions and sensorimotor skills. The ability to encode durations and use internal representations of time intervals is crucial for executing tasks effectively. Because a close link between the speed of information processing and subjective duration exists, as suggested by information-theoretic approaches, temporal coding, and timing behavior could reflect information processing efficiency (Grondin, 2010; van Wassenhove, 2023).

From the perspective of cognitive modeling, temporal perception is regarded as a mental ability supported by an *internal clock* known as the *pacemaker-accumulator model* (Gibbon, 1977; Matell & Meck, 2000; Treisman, 1963; van Rijn et al., 2014). This prominent theoretical framework is used to explain timing behavior across species (Matell & Meck, 2000). The pacemaker-accumulator model consists of two components: a pacemaker that generates rhythmic pulses and an accumulator that counts these pulses. The accumulated count encodes elapsed time and forms representations of time intervals, which are stored in memory to perform various timing-related tasks, such as temporal reproduction or discrimination (Treisman, 1963; Treisman et al., 1990). The internal clock model may be more suitable for *prospective timing*, which involves the continuous encoding of time, as opposed to *retrospective timing*, which encodes time as a chronological order of experienced events (Tsao et al., 2022).

Performing a time perception task involves temporal and non-temporal processes, implicating at least three cognitive functions: an internal clock, which encodes and transfers perceived time intervals into mental representations; a memory system, which stores the learned time intervals; and decision processes, which map the learned temporal information to a proper motor response. However, studies using time perception tasks often allocate the task effects to the internal clock alone, regardless of the possible influences from non-temporal processes, such as the decisional ones.

In the current study, we chose to use the cognitive architecture called Adaptive Control of Thought – Rational (ACT-R, Anderson et al., 2004; Anderson 2005; Taatgen et al., 2007; van Rijn et al., 2014) to recover the individual variability of both temporal and non-temporal processes in a duration change detection task. Temporal processes were modeled as a “ticking” internal clock and non-temporal processes were modeled as decision thresholds for the final output. We assumed both cognitive processes would contribute to the behavioral performance. Hence, we addressed three main questions: (1) Does sensory adaptation reduce, and novelty increase, the rate of the internal clock? (2) Does clock speed reflect the amount of cortical sensory processing? (3) Does motion distort time perception?

Temporal adaptation often occurs when individuals are exposed to repeating, unchanged, or stable stimuli, causing participants to perceive the duration of subsequent stimuli to be shorter than their actual duration. This is referred to as *temporal compression*. For instance, both short (340 ms) and long (860 ms) durations cause temporal compression after adaptation (Curran et al., 2016). When the comparison image is a repetition of the standard stimulus, its duration is judged to be shorter than that of the standard image. This effect has been observed for both face and non-face stimuli within the same paradigm (Matthews, 2015).

On the contrary, novel stimuli are often perceived to be longer than their actual duration, a phenomenon referred to as *temporal expansion* or *temporal dilation*. Researchers have found that novel stimuli, also known as oddballs, are perceived to last longer than they actually are, resulting in an expansion of subjective time (Matthews, 2011; Pariyadath & Eagleman, 2007; Tse et al., 2004; van Wassenhove et al., 2008). Sadeghi et al. (2011) reported both temporal compression of repeated stimuli and temporal expansion of novel stimuli in the same task. The temporal expansion caused by oddballs is robust across visual and auditory modalities (Birngruber et al., 2014; van Wassenhove et al., 2008), with its magnitude modulated by the amount of novelty, the likelihood of the oddball occurrence, and the serial position of the oddballs (Kim & McAuley, 2013).

Several working hypotheses could explain the contrasting effects of adaptation and novelty on time perception. One information-theoretic perspective suggests that temporal expansion results from increased demands for information processing caused by surprise (the oddballs). While enhanced arousal could account for the oddball effect, a study that manipulated arousal with salient and neutral oddball stimuli has ruled out the role of arousal in lengthening perceived time (Pariyadath & Eagleman, 2007), leaving the information-theoretic assumption as a more parsimonious option.

Neurophysiological studies have added more evidence to the information-theoretic approach of time perception by observing changes in sensory cortex simultaneous to temporal distortions. Sensory suppression due to the repeated presentation of the same stimuli (*i.e.,* neural adaptation) shows a reduced information processing rate (Kohn, 2007), which could account for the shortened perception of time. Meanwhile, sensory adaptation might enhance the sensitivity to novel stimuli, resulting in increased cortical excitability (Ranganath & Rainer, 2003) and a faster information processing rate when the upcoming stimuli are novel, causing longer subjective time (Eagleman & Pariyadath, 2009). In visual temporal judgment tasks, researchers have related a higher visual neural activity to a higher likelihood of reporting the stimulus to be longer (Noguchi & Kakigi, 2006).

Similarly, the middle temporal area of the monkey visual cortex has shown correlated activity to the degree of novelty, while the same experimental stimuli have induced a temporal distortion effect with human participants (Sadeghi et al., 2011). Note that neural adaptation might not reduce the coding efficiency of the repeated stimuli (van Wassenhove & Lecoutre, 2015). Instead, it is the noise that is attenuated during adaptation while the most informative parts are still encoded (Benda, 2021).

A direct contribution of the sensory cortex to the encoding of intervals was found in the study by Reinartz et al. (2024). Researchers trained two sets of rats to discriminate the duration and the intensity of vibrations respectively. Using optogenetic modulation of vibrissal somatosensory cortex, the authors found that the excitation dilated rats’ perceived duration and amplified their perceived intensity. These results indicate common neural substrates for temporal and sensory processes albeit with distinct coding mechanisms (Reinartz et al., 2024).

Intriguingly, motion perception seems to alter the time perception so that the internal clock speed might be susceptible to external speed, as evidenced by various studies comparing moving stimuli to static stimuli. For instance, the duration of moving targets is often overestimated, and the magnitude of temporal expansion increases with target speed (Brown, 1995; Kaneko & Murakami, 2009). Similarly, Kline & Reed (2013) have reported that a motion illusion elicited by a looming or a receding visual object leads to an overestimation of its duration compared to the static condition.

Cognitive theories propose that the expanded perception of intervals with speed could be attributed to the increased amount of experienced changes. Faster stimuli or moving objects are often associated with a higher level of perception and information processing, leading to longer subjective timing (Poynter, 1989).

However, some studies, such as Johnston et al. (2006), present a contrary finding concerning the impact of speed on time perception. In their study, perceptual adaptation to temporal frequencies at 5, 10, or 20 Hz caused participants to underestimate, rather than overestimate, the duration of subsequent test stimuli (always 10 Hz) appearing at the adapted spatial location. The authors demonstrated that the magnitude of the temporal compression was positively related to the speed of adaptors, with 20 Hz eliciting the strongest temporal compression (Johnston et al., 2006).

It noteworthy that methodological discrepancies might explain the divergent findings in the effect of motion speed on time perception. For instance, in Johnston et al. (2006), the adaptor served as a separate stimulus from the test stimuli that participants judged for duration. In contrast, Brown (1995) and Kline & Reed (2013) used the adaptors as the test stimuli. Meanwhile, the compression effect observed in Johnston et al. (2006) might be understood as the adaptation effect when the adaptor and test stimuli moved at the same frequency (both 10 Hz), analogous to a reference measure. In that case, the 5 Hz adaptor relatively reduced the adaptation effect and the 20 Hz adaptor relatively increased the adaptation effect.

In the current study, we examined three possible accounts linking timing and sensory processes using a duration change detection task of visual gratings. To test the role of adaptation, we fitted the clock of the cognitive model ACT-R with or without adaptation/novelty processes to each individual data, and tested whether clock groups differed in their perceived novelty. Our working hypothesis was that if clock adaptation is caused by sensory adaptation, participants whose clock does not respond to changes (unadapted clock) should yield reduced perceptual novelty effect compared to participants whose clock responds to both the adaptation and novelty processes (oddball clock). To further test the relatedness between timing and sensory processing, we decoded the sensory cortical features from EEG activity and examined if they were related to individuals’ clock speed. Our working hypothesis was that if clock speed reflects the amount of sensory decoding, sensory features should be correlated to the speed of the ACT-R clock. Last, to test motion speed tuning of timing, we tested whether individual clocks were modulated by the temporal frequencies of visual moving gratings. Our working hypothesis was that if visual motion modulates the clock, different temporal frequencies should yield different clock speeds.

## Results

### ACT-R models simulate the human behavior

Forty participants completed a temporal oddball task (Fig. 1a). They were required to detect odd durations that were either longer or shorter than the standard duration (+/- 20 %, 40 %, and 60 % of 600 ms). Participants responded by pressing the ’j’ key on the keyboard (Fig. 1b). The gratings could take four different temporal frequencies (motion speed, 0/5/10/20 Hz), which were counterbalanced within blocks (Fig. 1c).

**Fig. 1:**
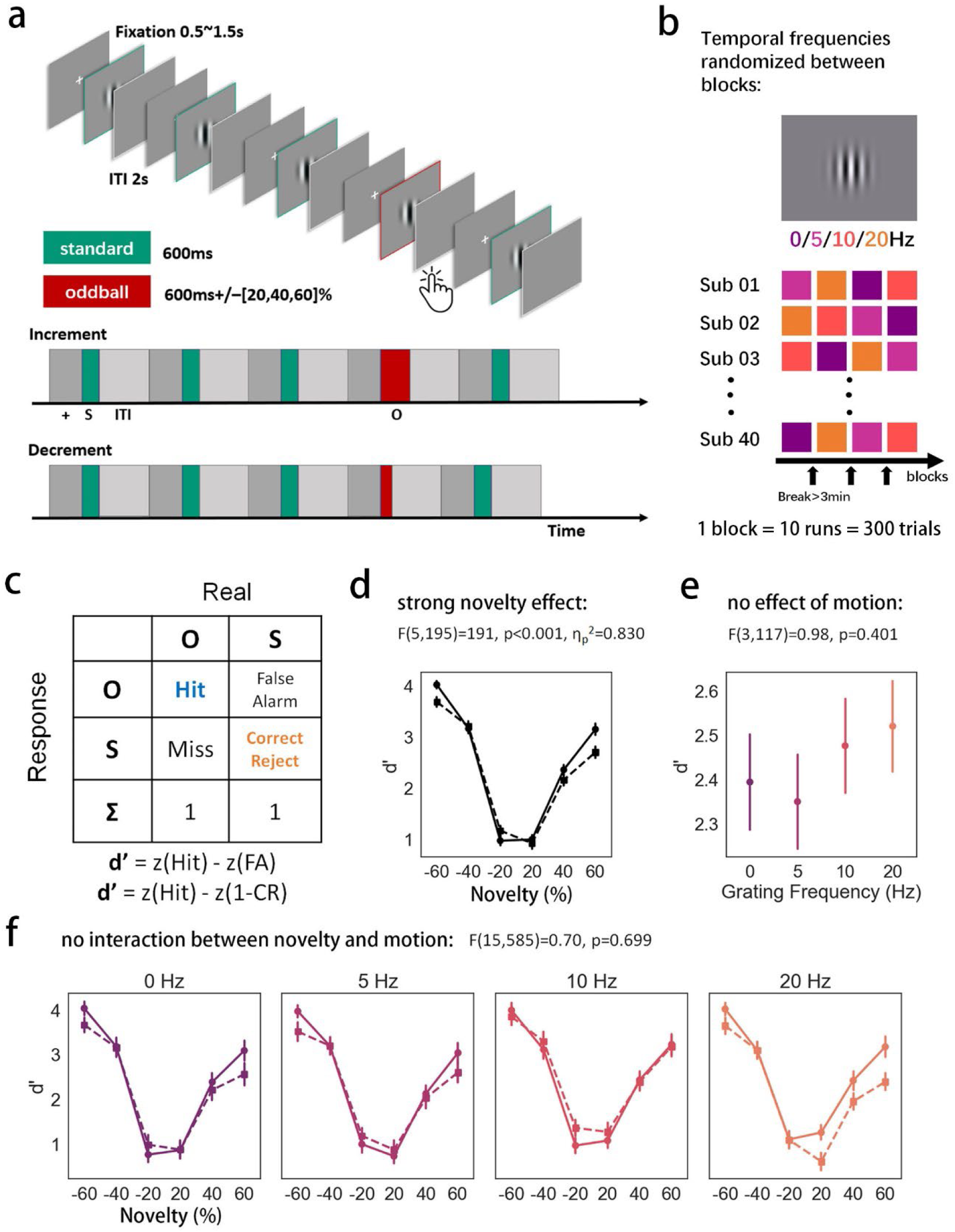
Experimental procedure as well as human and modeled performance. **a,** The trial sequence in the temporal oddball task. Each trial began with a fixation cross lasting between 0.5 and 1.5 s followed by a drifting (5, 10 or 20 Hz) or a static (0 Hz) grating. Most gratings (90%) appeared on screen for 600 ms, referred to as the standard. Targets (oddballs) appeared randomly (10%) in the trial sequence with either one of three levels of duration change as compared to the standard: 20%, 40%, and 60%. In the decrement block, the duration of oddball gratings was shorter than the standard ones (240 ms, 360 ms, 480 ms). In the increment block, the duration of oddball gratings was always longer than the standard ones (720 ms, 840 ms, 960 ms). Participants were asked to detect the oddball gratings (*i.e.*, changed durations) by pressing the “j” key during the inter-trial interval (ITI, 2 seconds) as a response to the detection (no response was required if no oddball was detected). The acronyms beneath the timeline denote fixation (+ in dark gray), standard stimuli (S in green), ITI (light gray), and oddball (O in red). **b,** Four temporal frequencies (0/5/10/20 Hz) alternated between blocks, and the order of temporal frequencies was randomized between participants. Participants took a break for at least three minutes between blocks. **c,** Behavioral metrics computed using the table of confusion. Hit rate (HR) indicates the accuracy in detecting targets (oddballs). Correct rejection rate (CR) indicates the accuracy in detecting the non-targets (standard). d’ is a synthesized score based on HR and CR. **d,** A strong effect of novelty intensity on human d’ was found. The detection sensitivity (d’) increases with relative oddball duration change. Solid line is human data. Dashed line is ACT-R simulated data. **e,** No effect of motion. **f,** No interaction between novelty and motion. Statistics in **d, e, f** indicate the experimental effect with human data. Error bars in **d, e, f** denote one standard error to the mean (SEM).

The original human performance as indicated by detection sensitivity (*d’*) clearly shows the novelty effect for the *d’* graded over novelty amount (Fig. 1d, f). To understand how the temporal and non-temporal processes work together to encode time intervals in this oddball paradigm, we designed an ACT-R model and fitted it to the individual behavioral data (Fig. 2a the general architecture, Fig. 2b the zoom-in temporal processes).

**Fig. 2:**
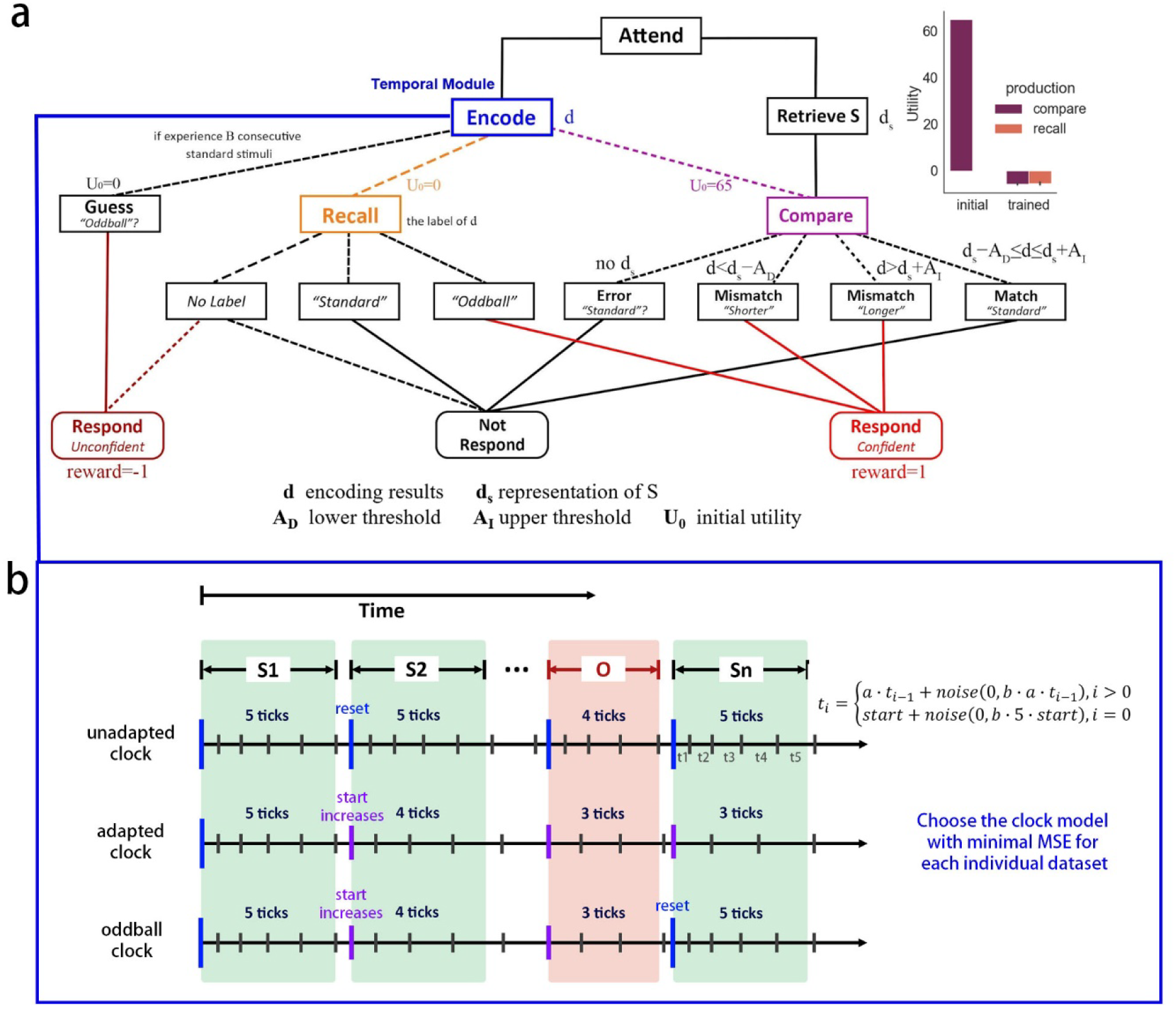
ACT-R design and clock models. **a,** ACT-R design to perform the temporal oddball task. **b,** Three internal clock models were designed depending on whether the clock responds to repetition or to novelty. S1 to Sn are standards (green). O is an oddball (red, shorter than standard in the given example). In the unadapted clock (top), the parameters of the temporal module are not affected by repetition or by novelty so that the number of ticks is nearly identical for the same durations (*e.g.,* always 5 ticks for the standards). The adapted clock (middle) adapts to repetition but does not respond to novelty. Thus, the number of ticks reduces for the same duration over trials (*e.g.,* ticks for standards changes from 5 to 3). The oddball clock (bottom) assumes both adaptation and novelty processes of the internal clock so that the number of ticks reduces for the same duration over trials in the absence of an oddball detection. When detecting an oddball, the clock resets and the number of ticks increases (*e.g.,* the number of ticks for standards decreases from 5 to 4 before an oddball detection, and “resets” to 5 after an oddball is detected).

Initially, the model attends to the stimulus on screen and starts encoding its duration in the temporal model (Fig. 2a). The encoded result (denoted as *d*) is compared to a learned representation of the standard duration (denoted as *d_s_*). If the difference between *d* and *d_s_* is lower than *A_D_* or higher than *A_I_* (the lower and upper thresholds, respectively), the model perceives the duration as an oddball and outputs ‘*j*’. Otherwise, the model perceives the duration as a standard. Herein, the decisional process serves as non-temporal processes for the final behavioral response.

Additionally, the utility variable decides the likelihood of the path to be selected. Initially, the model *encodes-compares* the stimuli for the response. After experiencing some stimuli, the model learns the labels and can identify stimulus type by recalling the label of *d*. This process is *encode-recall*.

Such a change can be shown by the change of utility before and after finishing the experiment by the model (Fig. 2a upper right corner). If the model responds using the comparison or the retrieved labels, a confident response is generated. If the model responds based on a guess, an unconfident response is given. A response with confidence gets a positive reward; a response without confidence gets a negative reward.

The *temporal module* was introduced to the ACT-R system by Taatgen et al. (2007) to model the internal clock based on the pacemaker-accumulator account of interval timing (Gibbon, 1977; Matell & Meck, 2000; Treisman, 1963; van Rijn et al., 2014). It consists of a pulse generator that produces ticks and an accumulator that counts these ticks. The interval between two ticks (*tick length*) increases as time progresses, akin to a metronome that ticks more slowly over time. This logarithmic growth of the encoded interval by the tick counts adheres to the *scalar property* of time and the Scalar Expectancy Theory (Gibbon 1977; Gibbon et al., 1984). The temporal module contains two main parameters, *start*, which decides the initial tick length, and *a,* which decides how fast the tick length increases (Fig. 2b).

Crucially, we confirmed the resemblance of task performance by the model to that by humans. The ACT-R simulated data captures the v-shaped *d’* changes over novelty amount, similar to human data (Fig. 1d, f).

### Novelty effect: clock adaptation differs from sensory adaptation

To study if the clock adaptation is caused by sensory adaptation, we designed three clock models based on findings we reviewed regarding how the internal clock reacts to repetition or novelty. Three models were fitted separately to each participant. The three tested clocks were (Fig. 2b): (i) The *unadapted clock*, which assumes that the parameters of the internal clock do not change over the trial sequence. In the unadapted clock, the internal clock always resets to its initial state whenever a stimulus appears regardless of whether the stimulus is a standard or an oddball (Fig. 2b, top); (ii) The *adapted clock*, which only responds to adaptational processes. The internal clock becomes slower over trials by increasing the parameter *start* in the temporal module every few trials. The clock adaptation process continues until the current run ends and resets in each run (Fig. 2b, middle); (iii) The *oddball clock*, which responds to both adaptation and novelty. That is, the parameter *start* increases every few standard trials. There are two cases when an oddball clock resets: at the beginning of each run and when detecting an oddball duration (Fig. 2b, bottom).

The best clock model for each individual dataset was decided by the minimal mean squared error (MSE) between the task metrics of the model vs. those of humans. With this approach, we found that the unadapted clock best explained 22 individual data whereas the oddball clock best explained the remaining 18 individual data (Fig. 3g). None of the individual data could be best explained by the adapted clock. This was not surprising since the adapted clock only models the clock adaption processes, while the current task involves both adaptation and novelty processes.

**Fig. 3:**
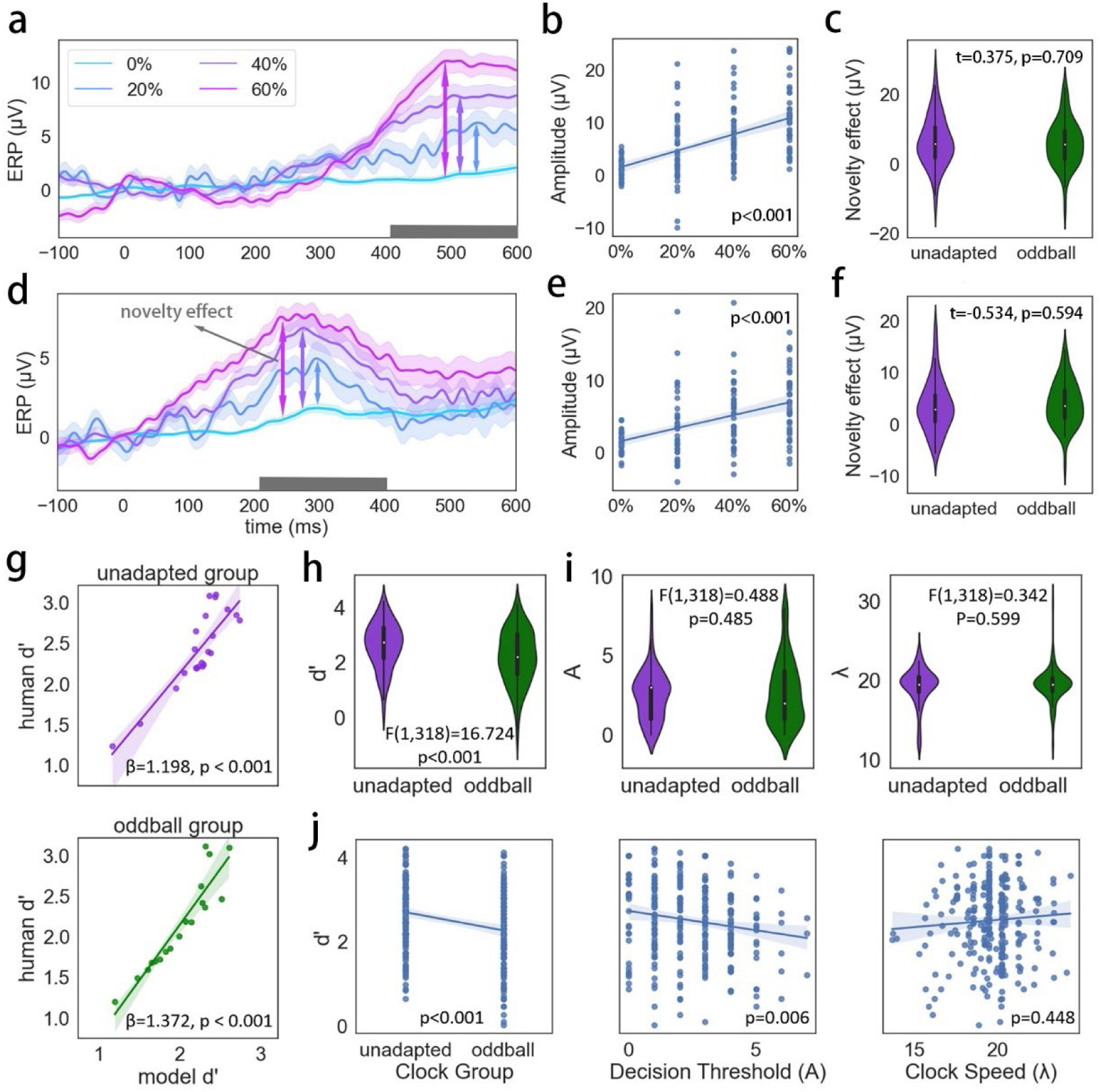
Cortical novelty effect with all grating frequencies. **a,** Offset ERP at Pz elicited by the standard stimuli (0 %) and the different levels of oddballs (20 %, 40 %, 60 %) in the decrement condition. Shaded lines denote one standard error to the mean (SEM). **b,** The mean ERP amplitude in the marked time window in **a** linearly increased with the percentage of duration change. **c,** there were no group differences for the novelty effect (as shown by non-significant one-way t-tests) between the unadapted (purple) and the oddball (green) clocks. **d, e, f,** Same analyses as **a, b, c** but with increment trials. The analysis was performed over all grating frequencies. **g,** We grouped participants according to their best-fitted internal clock models. The ACT-R performance resembled human behavior in both groups. **h,** Clock groups differed in their task performance. **i,** Clock groups did not differ in the decision threshold (A) or the clock speed (*λ*). **j,** The multivariate linear regression shows that both the clock type and decision threshold affected the behavioral outcome (human *d’*). However, the effect of clock speed was not significant. Translucent bands in **b, e, g, j** around the regression line denotes the 95% confidence interval for the regression estimate.

We then explored whether sensory adaptation could account for clock differences. If this was the case, clock groups would differ in their sensory responses to adaptation. Sensory adaptation is typically characterized by a larger amplitude of brain responses to the presentation of oddballs as compared to those elicited by the standards, known as the novelty effect (Amenedo & Escera, 2000; Benda 2021; Robinson et al., 2021). In time perception, the novelty effect is expected at the offset of the oddball (Ofir & Landau, 2022; van Wassenhove & Lecoutre 2015) as the change in duration would only be realized after the offset of the visual stimulus. For the analysis, we time-locked the EEG to the stimuli offsets and baseline-corrected using +/- 50 ms the stimulus offset (Ofir & Landau, 2022).

Our results showed that the amplitude of the ERPs increased linearly with greater duration changes at the midline EEG channels (Fig. 3, S1) in both decrement (Fig. 3a, b) and increment (Fig. 3 d, e) trials.

The sensory novelty effect was computed as the oddball responses minus the standard responses (double headed arrows in Fig 3a, d). We grouped participants according to their best-fitted model and found no differences in the novelty effect between the groups (Fig. 3c, f). This observation suggests that the sensory novelty effect may not reflect a clock process, refuting the earlier assumption. Hence, our results suggest that the EEG novelty effect in this oddball task may not account for the clock adaptation, and rather that clocking mechanisms for time perception may be fundamentally distinct from low-level sensory adaptation and change detection.

To further understand the possible differences between the distinct clock models, we analyzed the behavioral differences between clock groups. We found that the clock groups differed in task performance with the unadapted clock group showing a higher *d’* than the oddball clock group (Fig. 3h). We then pursued our analysis on the fitted temporal parameter (i.e. the clock speed *λ*) and the non-temporal parameters (i.e. the decision thresholds *A_D_*, *A_I_*). Our results showed that neither the clock speed nor the decisional threshold could account for the clock difference (Fig. 3i). Interestingly, running a multivariate linear regression model, we found that both the clock model and the decision threshold explain the *d’*, but not the clock speed (Fig. 3j). Altogether, these results stress the contribution of the non-temporal parts of the model to the final behavioral outcomes. This observation confirms the necessity of separating the non-temporal from the temporal explanatory factors when studying time perception.

### Decoded sensory features delineate sensory and clock processes

Having tested our first working hypothesis, which lead to the differentiation between clock and sensory adaptations, we then turned to our second working hypothesis and tested whether the clock speed reflects sensory processing amount. For this analysis, we decoded the neural features that could predict sensory processing. Our rationale was that if sensory and clock processes are related, decoded sensory features should be predictive of clock behavior.

We separately selected the 600 ms time window before the onset of the standard stimulus (henceforth, “stimulus-off” epochs) and the 600 ms time following the onset of the standard stimulus (henceforth, “stimulus-on” epochs). We only used the standard trials because we wanted to exclude motor evoked activity (mostly in response to oddball trials). Standard trials also provided 90 % of the collected EEG, providing a more reliable signal. We computed the power spectral density (PSD) separately for each time window using the Welch method (Fig. 4a). We further performed FOOOF to parameterize the raw PSDs and reconstruct the signal (Donoghue et al., 2020). The FOOOF parameterization gives an estimation of possible peaks as well as the aperiodic fit to the 1/f background. For each frequency band, if multiple peaks were identified, we took the peak with the highest power as the band feature (three features per peak – the peak frequency, the power, and the bandwidth). Meanwhile, we included the aperiodic parameters to the feature vector. This provided 18 features for each PSD (Fig. 4b).

**Fig. 4:**
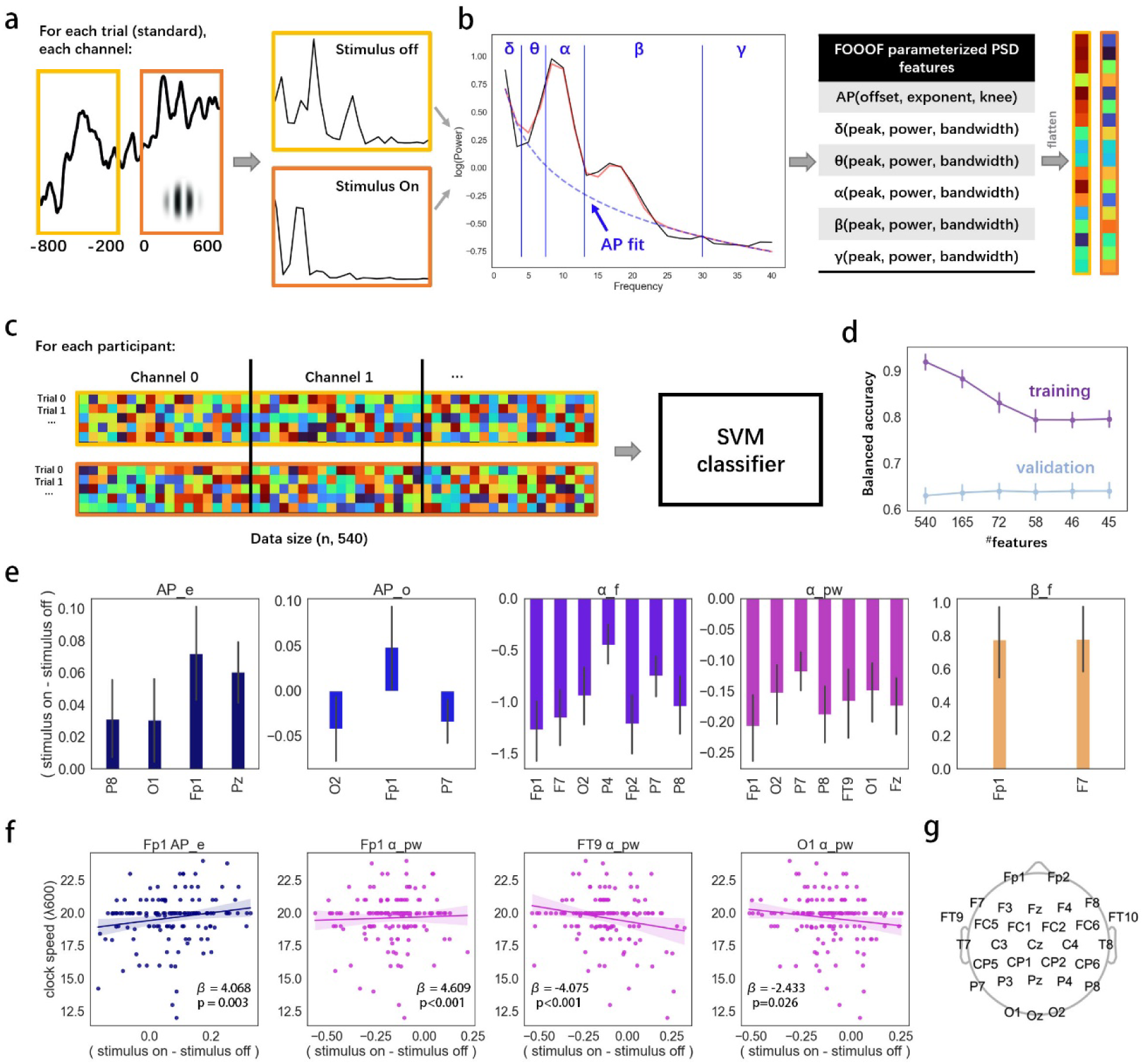
SVM classification and sensory features. **a,** Time window for stimulus-off and stimulus-on epochs. EEG epochs were transformed to welch-PSD (right panels). **b,** The raw PSDs were parameterized using FOOOF. We combined aperiodic and band parameters as feature vectors for the PSDs. **c,** Individual SVM classifiers were trained with features concatenated from all thirty electrodes to discriminate stimulus-on from stimulus-off data. **d,** Example of one feature selection session. Non-informative or redundant features were efficiently reduced over the modeling runs. The remaining 45 features maintained equivalent performance on the validation dataset compared to the initial model with 540 features. **e,** Features can linearly discriminate the sensory processing. **f,** Sensory features that linearly relate to the clock speed λ_600_ in standard trials. **g**, Layout of the thirty EEG channels. Error bars (shades) in **d, e, f** denote a 95% confidence interval. AP_e: exponent of the aperiodic activity; AP_o: offset of the aperiodic activity; α_f: alpha peak frequency; α_pw: alpha power; β_f: beta peak frequency.

We then trained individual SVM classifiers to discriminate between the PSDs of stimulus-off and stimulus-on epochs. Efficiently trained classifiers thereby highlighted the cortical features of sensory processing. For each trial, we concatenated PSD features from all 30 electrodes (Fig. 4g), which provided 540 features for either stimulus-off or stimulus-on epochs (Fig. 4c). In feature selection iterations, we dropped features that did not significantly affect the overall performance in each training, and subsequently obtained much fewer converging features (Fig. 4d). The balanced accuracy showed that the performance on the validation dataset remained stable with a reduced number of features, even though the performance on the training dataset was dropping. This indicated good control of the over-fitting of classifiers and of the decoding efficiency of the remaining features. Fig. 4d gives an example of one feature selection session. Overall, we ran 100 feature selection sessions. The final number of features converged to between 25 and 51 with an average of 34.31 ± 4.70. The converged performance on the validation dataset was 0.633 ± 0.003.

Because the convergence on the number of sensory features slightly differed between training sessions (Fig. S2), we ran an additional 100 feature selection sessions with the permutated data by shuffling the data labels. This analysis provided a feature probability distribution. We used the highest probability in permutation tests as the cutoff and selected 23 final sensory features (Fig. S3).

Although feature selection provided a clue on possible cortical features that contribute to the sensory processing of the standard durations, the SVM we used was nonetheless non-linear. To have a straightforward visualization of how the features changed with sensory processing, we performed t-tests of each of the 23 features between the stimulus-on and stimulus-off epochs. The results showed that all features could linearly inform sensory processing (Fig. 4e). The most prominent sensory features were: the exponent of the aperiodic activity (AP_e), which was steeper during sensory processing; the alpha peak frequency (α_f), which decreased during sensory processing; and the alpha band power (α_pw), which decreased during sensory processing, in agreement with the typical gating effect (Jensen & Mazaheri, 2010; Klimesch et al., 2007). Other sensory features that affected the SVM performance were the broadband offset of the aperiodic activity (AP_o) and the peak frequency of the beta band (β_f). Because the latter two features concerned less than 3 channels, we prefer not to draw firm conclusions from those observations.

Last, we analyzed the relationship between the internal clock and the observed features. Four features showed a significant linear relationship with the clock speed λ_600_ in a multivariate linear regression (adjusted R^2^=0.144). They were the exponent at Fp1 and the alpha power at Fp1, FT9, and O1 (Fig. 4f). A faster clock was observed with increasing exponent and alpha power at the prefrontal site and decreasing alpha power at the frontal-temporal and occipital sites. These observations suggest that the internal clock is related to the part of the cortical activity elicited by the presentation of the standard stimulus, thus suggesting that sensory processing of the standard duration is important for time estimation and internal clocking. However, because the novelty effect previously observed did not relate to the clock differences, sensory processing at large and clocking mechanisms are not identical.

### Motion affects sensory processing but is unrelated to the internal clock

Lastly, to address our last question pertaining to motion and its effect on the clock, we examined whether motion was related to early sensory processing or to the cognitive components of the internal clock. We already demonstrated by the task performance (Fig. 1e, f) that motion did not alter *d’*. Here, we further show that motion (in log scale) linearly related to the exponent of the aperiodic activity at P8 via the linear regression analysis (Fig. 5a). However, this sensory feature did not overlap with the sensory features that linearly related to the internal clock. Meanwhile, neither the decision threshold nor the internal clock seemed affected by motion speed (Fig. 5b,c). Together, the results provide no evidence that external speed can modulate internal speed.

**Fig. 5:**
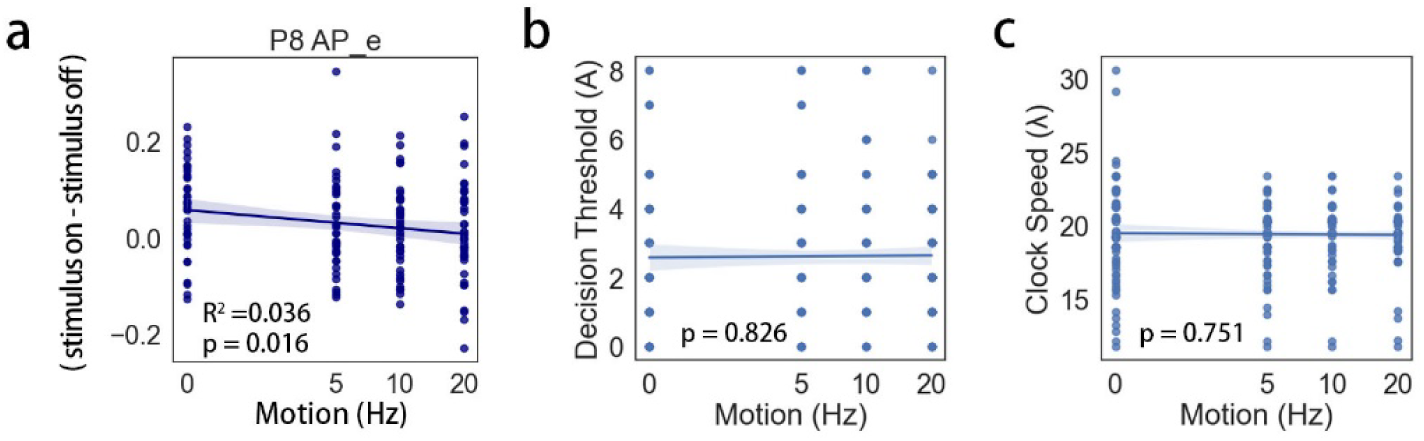
Motion effect on sensory processing and cognitive functions. **a,** Motion (grating frequency in log scale) linearly modulates the exponent of the aperiodic activity at P8 (one of the sensory features). b, Motion does not modulate the decision boundary A. **c,** Motion does not modulate the clock speed λ. Motion in all graphs is in log scale.

## Discussion

To disentangle the relationship between sensory processes and the internal clock, we examined three main claims that have been raised in the literature. The first claim is that sensory adaptation affects the internal clock (Curran et al., 2016; Birngruber et al., 2014; Eagleman & Pariyadath, 2009; Matthews, 2011; Matthews, 2015; Pariyadath & Eagleman, 2007; Sadeghi et al., 2011; Tse et al., 2014). The second claim is that sensory processing is manifested as subjective time (Eagleman & Pariyadath, 2009; Noguchi & Kakigi, 2006; Roseboom et al., 2022; Sadeghi et al., 2011; Sherman et al. 2022). The third claim is that motion affects the internal clock by adjusting sensory processing demands (Brown, 1995; Johnston et al., 2006; Kaneko & Murakami, 2009; Kline & Reed, 2013). Our results do not provide evidence in favor of the first and third claims, while partially supporting the second claim.

Our current results align well with previous findings highlighting the importance of decision processes in timing tasks, notably using the drift-diffusion models (DDM; Gibbon et al., 1984; Balcı & Simen, 2016; Ofir & Landau, 2022; Schumacher & Voss, 2023; Simen, 2014, 2016; Toso et al., 2021). Our results consolidate the proposal that the decisional stage should be considered in modeling timing tasks (Schumacher & Voss, 2023). Nevertheless, there are important differences to consider between the ACT-R decisional processes and the DDM decisional processes. DDM involves the evidence accumulation processes after the offset of a stimulus: once the accumulated evidence surpasses a threshold, a decision can be made. According to the DDM account for timing, a decision could be ready before the termination of the timed stimulus since the accumulation process might hit the threshold earlier than the stimulus offset (Balcı & Simen, 2016). Such an early-formed response happens specifically for longer durations in the context of shorter durations - as long as the timed interval surpasses the boundary, a longer decision can be made without fully encoding the duration (Kristofferson 1977). While the evidence accumulation process is not shown with the current ACT-R design for simplicity, some indirect evidence from the presented ERPs indicate the happening of evidence accumulation in the current task (Fig. 3a, d). The time window of the novelty effect was later in the decrement runs than in the increment runs. In increment runs, the early formation of an oddball detection could occur due to the sufficient amount of accumulated evidence before the end of the stimulus; this is not the case with the decrement runs.

One of our main findings is that the clock speed in the ACT-R model linearly changes with the alpha power (Fig. 4f). Alpha activity, especially when observed at the parietal-occipital region, has long been regarded as fundamental in the perceptual encoding of visual information (VanRullen & Macdonald, 2012; VanRullen 2016). Studies have shown that an improved visual discrimination performance is associated with a decreased alpha amplitude (Hanslmayr et al., 2005). Meanwhile, the inhibited alpha power might also indicate selective attention (Klimesch et al., 2007). Taking the periodic sampling role of the neural oscillations during sensory encoding (Busch & VanRullen, 2010), the change of alpha power with increasing sensory processing indicates the internal clock might be associated with an adjusted perceptual sampling amount. The current finding of augmented sensory processing with a faster internal clock partially supports that time perception has a functioning role in information processing.

The implication of alpha oscillations in timing, and alpha peak frequency in particular, has a long and winded history (Kononowicz & van Wassenhove, 2016; van Wassenhove et al., 2019). Interestingly, in perceptual awareness studies, correlational and causal approaches have linked alpha peak frequency to temporal sampling (Tarasi & Romei, 2023; vanRullen & Koch, 2003) and more recently to evidence accumulation (*e.g.,* Cuello et al. 2022). In perceptual processing, a higher alpha peak frequency supports the segregation of visual information whereas a lower alpha peak frequency supports the integration of visual information (e.g. Ronconi, et al., 2023; Samaha et al., 2015; Sharp et al 2022, Wutz, Melcher, & Samaha, 2018). Using causal neurostimulation techniques, a recent study by Mioni and colleagues (2020) has shown that the “speed” of alpha would linearly reflect the speed of the pacemaker in the internal clock. However, our current results did not replicate the sensory sampling rate with the alpha speed – we did not observe a significant modulation of the clock speed by the alpha frequency. However, the results showed a marginal relationship between the clock speed and alpha peak (β= 0.588, p=0.093). Future studies might increase the sample size to verify the results. On the other hand, a recent study exploring endogenous timing during a resting-state (no sensory stimulation) failed to find any evidence for the implication of the individual alpha peak frequency in duration estimation although other alpha parameters were essential (Azizi et al., 2023). Hence, the alpha frequency contribution to temporal resolution might depend on the experimental paradigm.

Despite the clocking differences across individuals, the oddball durations elicited a clear novelty effect in brain activity irrespective of clock group (Fig. 3). The cortical novelty effect is typically reported as being caused by sensory adaptation processes (Garrido et al., 2009; Malmierca et al., 2014; Benda 2021) but more recent evidence also hints towards a tight link between duration adaptation and decision processes (Hayashi et al., 2015; Hayashi et al., 2018; Protopapa et al., 2019). For instance, brain activity in the inferior parietal lobule (IPL) decreases with the repeated presentation of visual durations, indicating a neural adaptation effect to a timed interval (Hayashi et al., 2015). IPL has functional importance in sensorimotor integration and receives inputs from both cortical connections and subcortical structures (Clower et al., 2001). The role of IPL in duration adaptation indicates that duration adaptation engages high-level processes not reducible to the sensory encoding of duration. Additionally, the existence of duration-sensitive neural populations (Protopapa et al., 2019) have also been shown to be part of a larger network (Hayashi et al., 2018) and to be context-dependent (Kulashekhar et al, 2022). The processing of durations is indeed orchestrated in distributed neural networks with multiple structures involved, including the bilateral parietal cortex, the right inferior frontal gyrus, the medial frontal cortex, the right parietal cortex, and the striatum (Gouvêa et al., 2015; Hayashi et al., 2018). Altogether, prior neuroimaging findings relying on high spatial resolution highlight the contribution of a global network to timing estimation, in line with predictions from animal studies (Paton and Buonomano, 2018). Herein, the disentanglement of temporal processes with decision-making processes aligns with our current observations seeing that the modeled internal clock processes are related to very few sensory processing features in our data (2 out of 45), suggesting that timing processes in this duration change detection task engage activity well beyond the sensory analysis of stimuli.

In summary, our current findings indicate that the clocking mechanisms for time perception may be fundamentally distinct from low-level sensory adaptation and change detection. The relationship between the clock speed and the alpha power indicates that the internal clock (as modeled cognitively using ACT-R) may capture the sensory processing amount at the cortical level.

## Methods

### Participants

A total of forty participants were recruited for the experiment (23 females, aged 18-36 yrs, M+/-SD 24.00+/-4.72). All participants reported normal or corrected-to-normal vision. The research was conducted in accordance with the Declaration of Helsinki (2013) and was approved by the Research Ethics Committee of Neurospin, CEA (under the ethical protocol: CPP NeuroSpin CEA 100 049 2018). All participants signed written informed consents and were financially compensated for their participation in the two-hour experiment. All participants were included in the final data analysis.

### Experimental Procedure

The experiment involved four blocks of an oddball duration change detection task (Fig. 1).

The main stimuli used in the oddball task consisted of drifting gratings, which were presented at four different temporal frequencies: 0, 5, 10, and 20 Hz, with 0 Hz representing the static condition and higher frequencies indicating faster drifting speeds (Fig. 1c). The temporal frequencies of stimulus were counterbalanced within individuals and between blocks. Each oddball block consisted of 300 trials, presented in ten runs. All gratings used in the task had the same spatial frequencies (2c/cm). They were displayed in the center of the screen with a visual angle of 2.3°.

To minimize the motion aftereffects caused by the temporal frequency of previous stimuli, we designed the grating frequency as a block condition. Furthermore, participants were given a three-minute break between blocks. During this rest time, a countdown timer starting from the 180s was provided.

Fig. 1b illustrates exemplar trials. Each trial began with a fixation cross displayed at the center of the screen for a duration randomly drawn from a uniform distribution ranging between 0.5 and 1.5 s. A grating was then presented on the screen for 600 ms in standard trials, which made up 90% of the trials in the task. In oddball trials, the duration of the gratings could deviate away from the standard duration (600 ms) by 20%, 40%, or 60%. In the decrement run, oddball stimuli lasted for 240 ms, 360 ms, or 480 ms, whereas in the increment run, oddball stimuli lasted for 720 ms, 840 ms, and 960 ms. Participants were required to detect these oddball durations and respond by pressing the “j” key on the keyboard during the inter-trial interval (ITI) of 2 seconds. Each run consisted of thirty trials, with 27 standard gratings and 3 oddball gratings (+/- 20%, +/- 40%, and +/- 60% each with the sign depending on the run).

Importantly, two successive oddball trials were separated by at least three standard trials. The five initial trials of each run always consisted of standard gratings. The decrement and increment runs were counter-balanced and alternated throughout the block.

### Behavioral Analysis

We evaluated human performance in the study using two metrics: the hit rate (HR) and the correct rejection rate (CR). HR represents the accuracy of detecting positive cases, while CR measures the accuracy of detecting negative cases.

We further computed *d’* as a synthesized score using the equation: *d’ = z(HR) – z(1-CR)*, where *z*() represents the z-score. However, it is worth noting that in some cases, HR and CR might approach 1, potentially leading to infinite z-scores. To address this, we adjusted HR and CR before applying the z-transform, following the method proposed by Hautus (1995). The adjusted HR and CR were then used to compute *d’*:

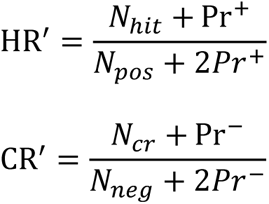

Where *N_hit_* is the number of hits; N_cr_ is the number of correct rejections. *N_pos_* is the total number of positive trials (oddballs); *N_neg_*is the total number of negative trials (standard). The *Pr^+^* and *Pr^-^* indicate the probability of positive and negative trials respectively. In the results, HR and CR refer to the original HR, CR while *d’* is calculated from their adjusted values (HR’, CR’).

### ACT-R Design

The Adaptive Control of Thought – Rational (ACT-R, http://act-r.psy.cmu.edu/) is a cognitive architecture used for simulating and understanding human behavior and cognition. ACT-R’s constructs are based on findings from experimental psychology. Researchers can customize ACT-R models to suit their assumptions about specific tasks, allowing for the study of various research questions related to attention, perception, memory, learning, and more, across a wide range of complexities.

ACT-R consists of several modules: the vision and audition modules model the sensory input processes. The motor module generates outputs that model the behavioral response in a cognitive task. There are two key modules in ACT-R: the *declarative module* and the *procedural module*, which are both essential to almost all ACT-R models. The declarative module models the declarative memory system where the learned information is stored in a form called *chunk*. The procedural module stores the procedural knowledge which are *if-then* statements defined by the user. When running ACT-R models, the conditional part (‘*if*’) of the production will be checked, and the production with the matched *if* will be selected and fired. If there are multiple productions that meet the selection criterion, the firing likelihood of the production is decided by its *utility*, which has an initial value and can be *reinforced* during the modeling. Similarly, each chunk in the declarative module has its *activation* value. The activation value decides the likelihood of the chunk being retrieved when multiple chunks meet the retrieval criteria and will decay or get reinforced based on the frequency and recency rule in ACT-R (Anderson et al., 2004; Anderson 2005).

#### ACT-R Architecture for the Time Oddball Task

The ACT-R Architecture designed for the time oddball task contained the basic cognitive components generally used in time perception tasks: time, memory, and decision (Matell & Meck, 2000; Parayadath & Eagleman, 2007). The individual parameters to be fitted are listed in Table 1. Best parameters were those that gave the minimal mean squared error (MSE) during the fitting:

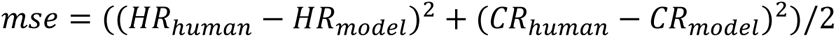

**Table 1.** Individual Parameters to Tune.

|  |  | Parameter | Interpretation | Search Range | Notes |
| --- | --- | --- | --- | --- | --- |
| Non-clock parameters | | $A_D$ | Lower threshold | [0,8] | Fitted to each grating frequency of decrement runs |
| | | $A_I$ | Upper threshold | [0,8] | Fitted to each grating frequency of increment runs |
| | | $B$ | Take a guess every B standard stimulus | [5,16] | Fitted to the whole task |
| Clock parameters | Adapted clock | $n_{adapted}$ | Start pulse increases every n trials in the adapted clock | [1,30] | |
| | Oddball clock | $n_{oddball}$ | Start pulse increases every n trials in the oddball clock | [1,30] | |
| | | $a$ | Tick dilating speed (time-mult) | [1.01, 1.29] | |
|  | All clocks<br>(but fitted<br>separately) | **start** | Starting tick length (start-<br>increment) | [0.006, 0.015] | Fitted to each<br>grating<br>frequency |

##### Encode-Compare

An ACT-R model was tailored to perform the time oddball task (Fig. 2a). The model attended to and the temporal module encoded the duration of the stimuli, with the encoding results denoted as *d*. Simultaneously, it attempted to retrieve a representation of the standard duration, represented as *d_s_*. Two individual parameters were fitted in the comparator: *A_D_* and *A_I_*, serving as lower and upper thresholds for decision-making. Specifically, the model judged a duration as an oddball when it falls above the upper boundary (*d_s_*+*A_I_*) or below the lower boundary (*d_s_*-*A_D_*).

Otherwise, it categorized the duration as standard. An exception occurred at the beginning of the experiment, where the model did not yet form a representation of the standard duration, so it encoded the first stimuli as standard, aligning with the experimental setup where the first five trials of each block were consistently standard. Following the chunk harvest mechanism in ACT-R, each encoded duration *d* would be moved to the declarative module to form the representations of oddball or standard durations once the trial was finished.

##### Encode-Recall

As the model encountered and learned a few oddballs, it became capable of identifying an oddball without relying on the comparator. Instead, it might attempt to retrieve the labels of the encoded durations and provide the corresponding response if the label was successfully retrieved. If the model failed to retrieve the label, it would guess regarding whether the duration was a standard or an oddball. Initially, the probabilities of both guesses were set to 50% due to equal starting utilities.

##### Encode-Guess

This part of the model incorporated the experimenter’s observation that participants, during the task, tended to anticipate a higher likelihood of encountering an oddball stimulus if they had been perceiving several standard stimuli consecutively. This behavior was captured in the “encode-guess” pathway, where participants were more likely to guess that the upcoming stimuli would be an oddball after perceiving a certain number of consecutive standard trials (denoted as *B* and fit for each dataset).

The responses provided by the model based on the comparator output or successful retrieval of labels were considered confident responses and were rewarded positively. On the other hand, responses derived from guessing were penalized with negative rewards. No response received no reward. Initially, the encode-compare utility was set to 65 and encode-recall was set to 0 so the model could well learn the representations at the beginning. At the end of the experiment, the utility of encode-compare and encode-recall became almost equal (Fig. 2a right upper corner), indicating a half chance of directly retrieving the label instead of mechanically comparing.

#### ACT-R Temporal Module

The *temporal module* was introduced to the ACT-R system by Taatgen et al. (2007) to model the internal clock based on the pacemaker-accumulator account of interval timing (Gibbon, 1977; Matell & Meck, 2000; Treisman, 1963; van Rijn et al., 2014). It consists of a pulse generator that produces ticks and an accumulator that counts these ticks. The interval between two ticks (*tick length*) increases as time progresses, akin to a metronome that ticks more slowly over time. This logarithmic growth of the encoded interval by the tick counts adheres to the *scalar property* of time and the Scalar Expectancy Theory (Gibbon 1977; Gibbon et al., 1984).

Although the temporal module operates independently from other modules in generating and counting ticks, ACT-R models can be designed to investigate the interaction between cognitive modules and the temporal module. The resulting count from the temporal module can be queried and accessed by other modules. Additionally, the timed interval in the temporal module is used by the declarative module to form a learned representation of the interval.

The ACT-R temporal module has been shown to successfully model human data in several *explicit timing* tasks studying either time perception (such as the time reproduction task by Rakitin (1998), or the bisection task by Penney et al. (2000)), or the interaction between timing and cognition such as the study by Zakay (1993). The model has also predicted human performance in a dual-tasking paradigm, when the interval needs to be *implicitly* coded for the successful completion of a non-timing task (Taatgen et al., 2007), achieving overall good simulation.

The temporal module encodes duration with ticks. The length (duration) of each tick is:

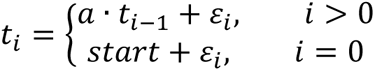

in which the noise *ε* is generated from the logistic distribution (approximation of a normal distribution) with 0 mean and a given *s*:

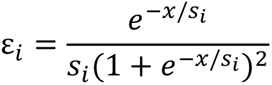

The *s* value is related to the variance of the distribution: *σ^2^=π^2^*s^2^/3*. In the temporal module *s* is computed by the equation:

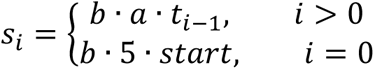

In this study, we kept *b* to its default value of 0.015 and we fit *a* and *start* to each block (grating frequency) for each participant. This is because both *start* and *a* determine the ticking speed (thus, the tick length) of the internal clock as can be told by the equations (*start* sets the initial tick length and *a* decides how fast the tick length increases).

#### Clock Adaptation

Following assumptions regarding whether the internal clock reacts to repetition or novelty, we designed three internal clock models using ACT-R, and fit them separately for each participant (Fig. 2b). We called these models the unadapted clock, the adapted clock, and the oddball clock. We describe their distinctive properties as follows:

***Unadapted Clock*** assumes no effect from either the adaptation or the novelty processes, positing that the temporal module always resets to its initial state in each trial.

***Adapted Clock*** assumes that the internal clock is subject to the adaptational process where its speed slows down gradually with time. This is achieved by lengthening the initial tick (*start*) every few trials for both standard and oddball trials.

***Oddball Clock*** adapts to repetition by lengthening the initial tick (*start*) every few trials, but such modulation occurs only for standard trials. Any detection of oddballs will reset the temporal module to its initial state, simulating the effect of novelty that increases the internal clock.

To avoid an overly rapid dilation of the initial ticks, the increase of *start* usually does not happen between two consecutive trials. Instead, it is controlled by a parameter denoted as *n_adapted* and *n_oddball*, indicating the number of preceding trials before *start* increases once in the adapted clock and oddball clock respectively (Table 1). The increment of start is set to 0.001 for both clocks for all participants.

#### Fitting Procedure

For each participant, we trained the three ACT-R models to fit the behavioral data: the unadapted clock, the adapted clock, and the oddball clock. The best fit was decided by the minimal mean squared error (MSE) of the ACT-R performance as compared to the human behavioral metrics (Hit Rates (HR) and Correct Rejections (CR)).

The fitting began with the unadapted clock and included four steps: (1) We fitted an initial decision boundary *A_I_* and *A_D_* to the increment and decrement trials respectively. (2) The clock parameters *a*, *start* were fitted to each grating frequency. (3) *A_I_* and *A_D_* were fine-tuned for each grating frequency. (4) Finally, we fitted *B* to the entire dataset.

When fitting the adapted and oddball clocks, the ACT-R model first inherited the *A_I_*, *A_D_*, *a*, *start* from the unadapted model and an initial *n_adapted* or *n_oddball* was fitted to the whole dataset. Then we re-fitted other parameters following steps (1)-(4) same as fitting the unadapted clock. Lastly, we re-fitted the *n_adapted* or *n_oddball* to the whole task.

#### The Clock Speed

We estimated the internal clock speed at each grating frequency per individual with the fitted ACT-R temporal module parameters.

A duration *D* can be represented as the summed length of n ticks:

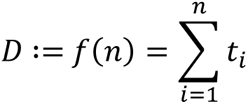

The inverse solution gives the number of ticks that encode duration D:

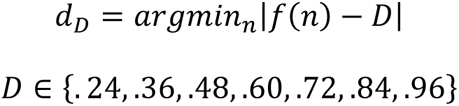

Where *d_D_* denotes the internal representation of duration *D* in the form of the number of ticks. To simplify the calculation, we omitted the noise term *ε* when estimating the *d_D_*, so each *t_i_* is estimated as:

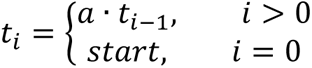

The average *d_D_* in one block is denoted by *λ*, which indicates an individual’s internal clock speed in this block. It is easy to infer that a higher *λ* indicates a faster internal clock for the block. Specifically, we calculated *λ_600_* with standard trials only (*D*=0.60).

### EEG Recording and Analysis

#### EEG Recording

Continuous EEG was recorded while participants were performing the oddball task. The Brain Products (GmbH, Germany) LiveAmp 32-channel system and actiCAP slim with gel-based active electrodes were used for high-quality EEG recordings. The impedance was kept below 25kΩ during the initial set-up and was checked by the experimenter during breaks. The layout of the actiCAP follows the International 10-10 System with FCz serving as an online reference. The online sampling rate was 500 Hz.

#### Preprocessing

EEG data were preprocessed using the EEGlab toolbox (v2022.0, Delorme & Makeig, 2004) in Matlab R2022b. The data were first re-referenced to the averaged mastoids (TP9, TP10) and then band-pass filtered within the range of 0.1 and 42 Hz. Continuous EEG was segmented into epochs, time-locked to the onset of the grating, including a period of 0.8 seconds before and 1.7 seconds after the onset. Artifacts arising from ocular movements (e.g., blinks, saccades), muscle activities, and disconnected channels were identified and removed based on the ICA results. Subsequently, the epochs were manually inspected to exclude trials with excessive artifacts that were not effectively removed by the ICA procedure. On average, 2.8% of the trials were excluded from each dataset.

#### Event-Related Potential (ERP)

The sensory adaptation was examined by the novelty effect evoked by oddball trials. EEG epochs after preprocessing were time-locked at the stimulus offset and re-aligned at the time window with 50 ms before and 50 ms after the offset (Ofir & Landau, 2022). ERPs were obtained by averaging EEG epochs over trials per individual.

#### Power Spectral Density (PSD)

PSD was performed on the standard trials (90% of all trials). For each trial, we took the 800 ms to 200 ms preceding the grating onset (the fixation period) as the stimulus-off epoch, and 0 ms to 600 ms following the grating onset as the stimulus-on epoch. We computed the PSD for each epoch using the Matlab function *pwelch()* .

### Feature Search with Support Vector Machine (SVM)

#### PSD Features

Raw PSD was parameterized and reconstructed with the FOOOF algorithm (Donoghue et al., 2020). The FOOOF parameterization gives an aperiodic fit as well as spectral peak fits (Fig. 3b). For the band features, we took the peak with the largest power within the frequency range of that band. Each band has three features: peak frequency, power, and bandwidth (denoted as *f*, *pw,* and *w* in the results). Five bands were defined: delta band 0.5-4 Hz, theta band 4-7.5 Hz, alpha band 7.5-13 Hz, beta band 13-30 Hz, and gamma band 30-40 Hz. Missing peaks at the certain band were denoted as zeros in the feature matrix and later were interpolated by feature mean during machine learning.

Aperiodic activity *L* was fitted using the equation (Donoghue et al., 2020):

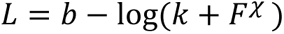

Where *b* is the broadband offset, *χ* is the exponent and *k* is the ‘knee’ parameter. In the current study, *k*=0 for all fittings is equivalent to fitting the aperiodic activity in log-log spacing.

Thus, each PSD was transformed into a vector of 18 features.

For each trial, we concatenated the 18 features from all thirty channels (Fig. 4g) and obtained 540 features for the classification (Fig. 4c).

#### SVM Training and Validation

SVM classifiers were trained with the Scikit-learn package on Python. We trained non-linear SVM with the Radial Basis Function (RBF) as the kernel. The kernel coefficient *gamma* is set to be 1/(n_features * X.var()). The regularization parameter *C* was chosen between [1e-5, 1e-4, 1e-3, 1e-2, 1e-1, 1, 10, 100] with the best fit during the cross-validation processes. We used five-split-cross-validation in the current study. The performance was measured using the balanced accuracy:

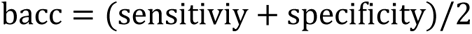

where sensitivity and specificity refer to the accuracy in detecting positive and negative cases respectively. In the current study, positive cases refer to stimulus-on PSDs, and negative cases refer to stimulus-off PSDs.

#### SVM Feature Selection

We computed the effect of each feature on the trained classifiers by muting them (*i.e.,* set the feature value to *nan*) sequentially, and observed the change in overall performance, the score was calculated as:

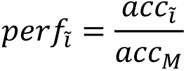

where 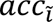 refers to the classifier’s accuracy when the i-th feature is muted and *acc_M_* refers to the original performance of the trained classifier. It is not hard to understand that if 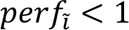, then removing the i-th feature would lower the model’s performance. In other words, the i-th feature is efficient and informative in the trained classifier. We thus trimmed redundant features and kept those features that affect overall modeling performance in the next modeling run and repeated the above feature selection step until the feature number converged (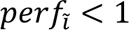 for all *i*), meaning no more redundant features in the classifier.

The feature selection modeling was repeated over 100 sessions to obtain the occurrence probability of each feature. We ran another 100 sessions with the permuted data by shuffling the data labels to form the permutated feature probability. We took the maximal feature probability in permutation as the cutoff for the sensory features (S3).

#### Sensory Features

SVM feature selection processes gave features that indicate sensory processing intrinsically in a non-linear way. We further examined if those features could linearly discriminate the sensory processing by looking for features with significant differences between stimulus-on vs. stimulus-off. Statistics were performed with t-tests.

### Statistical Analysis

Statistical analyses such as ANOVA, t-tests and linear regressions were performed with the Pingouin package on Python (Vallat, 2018).

#### Multivariate Outliers

Multivariate linear regressions were performed to assess the relationship between the sensory EEG features and the clock speed. We used the Mahalanobis distance to identify the outliers in a multivariate space (Leys et al., 2018), which is defined as:

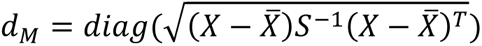

where X is an N × K data matrix with N observations and K dimensions (variates) and 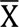 is defined as 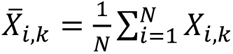 so that each column of 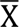 is the column mean of X. S is a K × K covariance matrix.

The *diag()* operation takes the diagonal of the matrix. This gives a 1 × N vector that d_M(i)_ is the Mahalanobis distance for observation X_i_.

A data point is considered an outlier if its Mahalanobis distance d_M_ meets the condition (Leys et al., 2018):

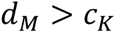

where 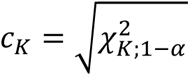 , indicating the square root of the upper-α quantile of the chi-squared distribution with K degrees of freedom, taking α=0.05 currently. Outliers were removed when fitting the multivariate linear regressions.

#### Multivariate Linear Regression

Similar to training SVM classifiers, we trained the model with all variables first and dropped variables one by one to observe the change of the adjusted R^2^, with reduced R^2^ indicating an informative variable. We kept the informative variables and re-trained the regression model until the adjusted R^2^ stopped increasing.

## Data and Code Availability

All data are available via G-Node at https://gin.g-node.org/cyj.sci/Time_Oddball. All code is available via GitHub at https://github.com/christina109/Time_Oddball.

## Acknowledgements

We thank J. P. Borst, T.W. Kononowicz, S.K. Herbst, M. Logie, and C. Grasso for their helpful comments on an earlier version of the manuscript. This work was supported by the EXPERIENCE Project of the European Commission H2020 Framework Program Grant No. 101017727 (to V.vW.).

## CRediT Author Statement

C.Y.J. contributed to conceptualization, methodology, software, formal analysis, investigation, data curation, writing – original draft, writing – review & editing, and visualization. A.R. contributed to conceptualization, methodology, and investigation. R.B contributed to investigation. V.vW. contributed to conceptualization, methodology, resources, writing – review & editing, supervision, project administration, and funding acquisition.

## Supplementary information

**Fig. S1:**
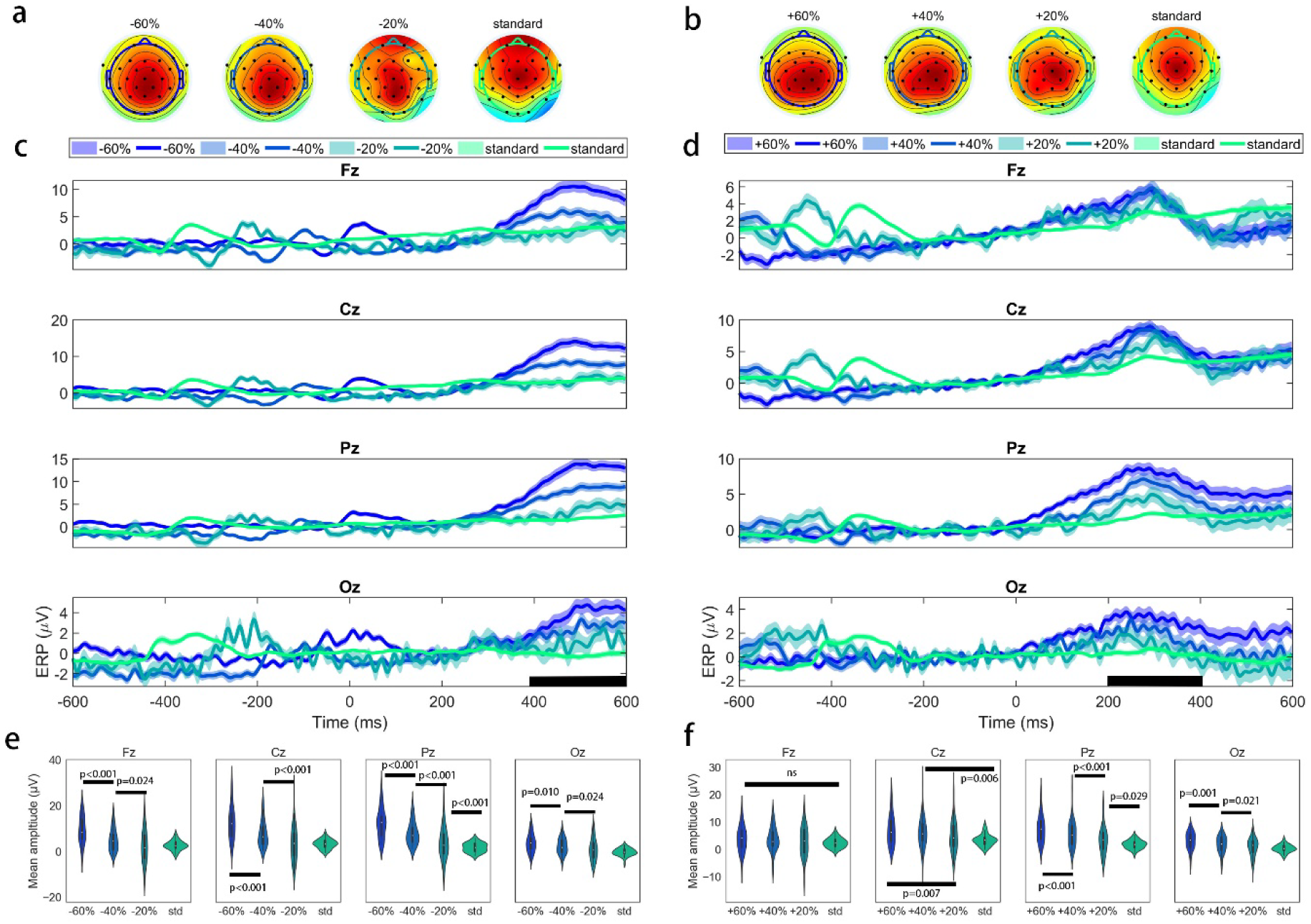
Novelty effect was prominent at the midline EEG channels. **a,** Topoplot of EEG between 400-600ms (marked time window in **c**) for the decrement runs. **b,** Topoplot of EEG between 200-400ms (marked time window in **d**) for the increment runs. **c, d,** ERP time-locked to the offset of the gratings with baseline corrected at -200-0ms at Fz, Cz, Pz, Oz (midline channels). The black line on the timeline indicates the time window where the strongest novelty effect was observed. The corresponding statistics are summarized in **e, f.** P-values in **e,f** indicate significance for pairwise comparisons corrected by Benjamini-Hochberg FDR method. *ns* indicates no difference between conditions.

**Fig. S2:**
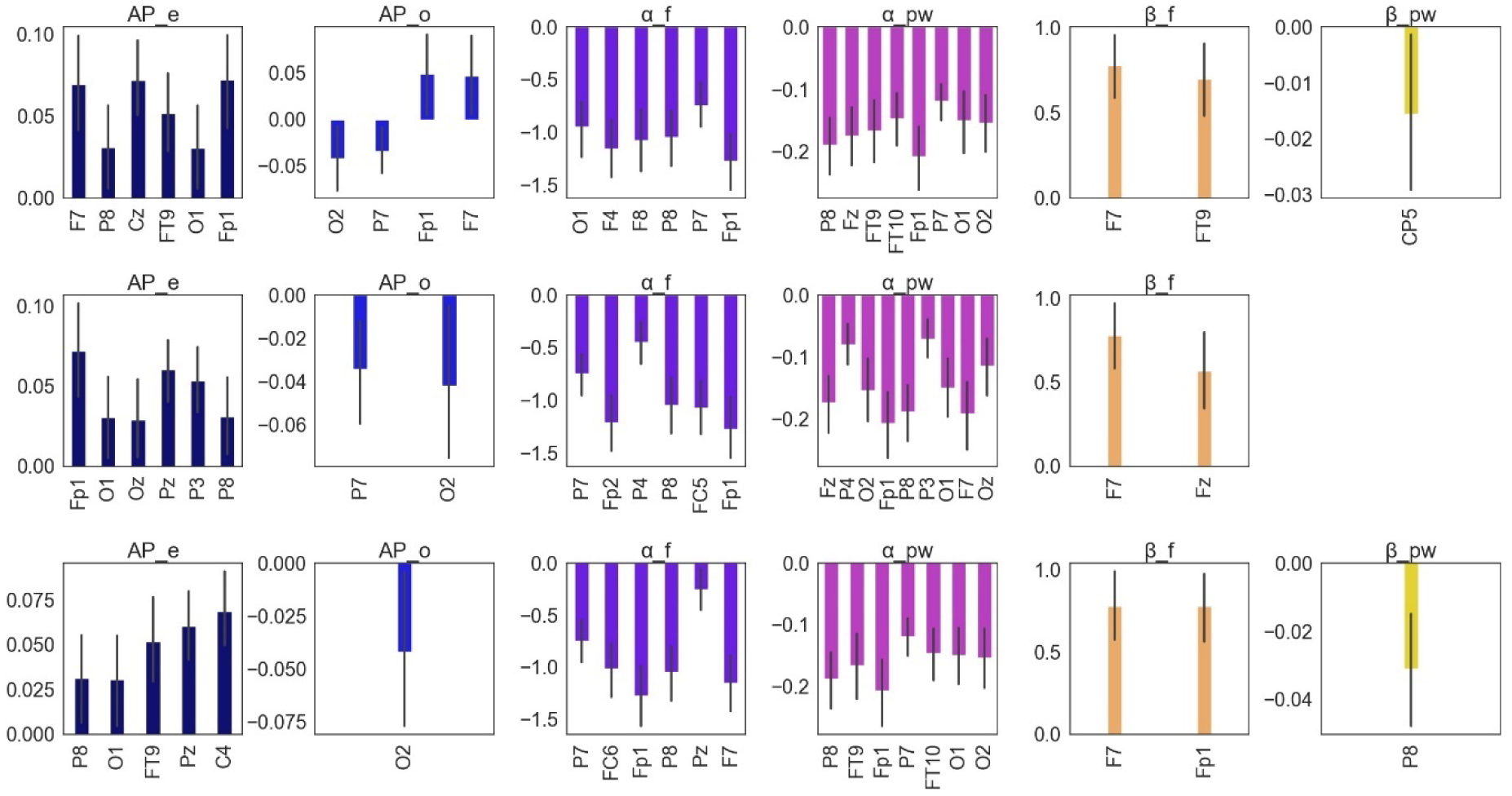
Sensory features slightly differ between modeling sessions. The graph shows the feature selection outcomes from three random modeling sessions (from top to bottom, sessions #3, #8, #12, respectively). These illustrate that even though the overall categories remained stable, the selected observational locations by model (e.g., the EEG channels) might differ across training sessions. This is easier to understand considering the EEG signals are correlated between adjacent channels.

**Fig. S3:**
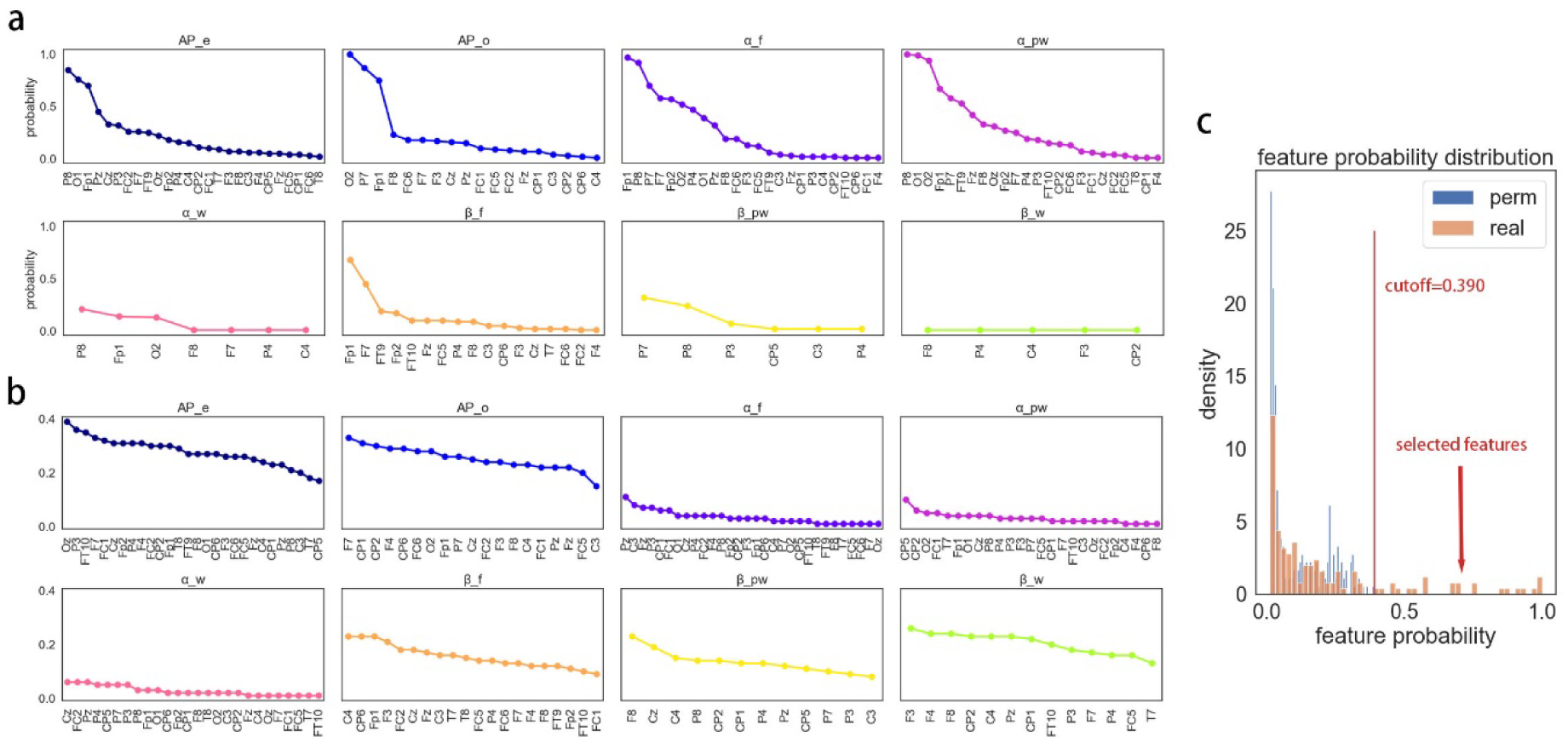
Probability of the selected features. a, Probability of the features with real data. b, Probability of the features with permutated data. c, The feature probability distribution for real and permutated data. The cutoff was the highest probability achieved during the permutation tests. We selected the sensory features with probability above the cutoff.

## Notes

https://gin.g-node.org/cyj.sci/Time_Oddball

https://github.com/christina109/Time_Oddball

### Competing Interest Statement

The authors have declared no competing interest.

